# Transcriptional regulation of voltage-gated sodium channels contributes to GM-CSF induced pain

**DOI:** 10.1101/386698

**Authors:** Fan Zhang, Yiying Wang, Dandan Zhang, Xizhenzi Fan, Hao Han, Xiaona Du, Nikita Gamper, Hailin Zhang

## Abstract

Granolocyte-macrophage colony stimulating factor (GM-CSF) induces production of granulocyte and macrophage populations from the hematopoietic progenitor cells; it is one of the most common growth factors in the blood. GM-CSF is also involved in bone cancer pain development by regulating tumor-nerve interactions, remodeling of peripheral nerves and sensitization of damage-sensing (nociceptive) nerves. However, the precise mechanism for GM-CSF-dependent pain is unclear. In this study, we found that GM-CSF is highly expressed in human malignant osteosarcoma. Rats implanted with bone cancer cells develop mechanical and thermal hyperalgesia but antagonizing GM-CSF in these animals significantly reduced such hypersensitivity. Nociceptor-specific voltage gated Na^+^ channels Nav1.7, Nav1.8 and Nav1.9 were found to be selectively up-regulated in rat DRG neurons treated with GM-CSF, which resulted in enhanced excitability. GM-CSF activated the Jak2 and Stat3 signaling pathway which promoted the transcription of Nav1.7-1.9 in DRG neurons. Accordingly targeted knocking down either Nav1.7-1.9 or Jak2/Stat3 in DRG neurons alleviated the hyperalgesia in rats. Our findings describe a new bone cancer pain mechanism and provide a new insight into the physiological and pathological functions of GM-CSF.

## Introduction

Granolocyte-macrophage colony stimulating factor (GM-CSF, or CSF2) was originally identified as a colony stimulating factor because of its ability to induce granulocyte and macrophage populations from precursor cells. It is one of the most common growth factors in the blood system (Croxford, et al.,2015). GM-CSF is also abundantly secreted by some tumor cells and plays a key role in regulating tumor-nerve interactions, remodeling of peripheral nerves and sensitization of damage-sensing nerves (nociceptors) by acting at receptors (Schweizerhof, et al.,2009). Apart of the bone metastases pain, GM–CSF was also shown to be involved in inflammatory pain, arthritic and neuropathic pains (Cook, et al.,2013; Cook, et al.,2012; Nicol, et al.,2018). In line with this, treatment with therapeutic monoclonal antibody against GM-CSF significantly suppresses arthritic pain, neuropathic pain and bone metastases pain (Cook, et al.,2013; Cook, et al.,2012; Nicol, et al.,2018). A recent study show that GM-CSF signaling contributes to pain-associated behavior that is independent of a gliosis and/or astrocyte response, suggesting that GM-CSF may directly activate sensory neurons (Nicol, et al.,2018). However, the precise mechanism for GM-CSF-dependent pain is unclear.

Dorsal root ganglion (DRG) neurons are the peripheral somatosensory neurons, and are the primary nerve cells responsible for nociceptive signal initiation and propagation. Receptor for GM-CSF (GM-CSFR) are found to be expressed in DRG and in peripheral nerves dispersed in the periosteum of mice (Schweizerhof, et al.,2009). Signaling cascades and mechanisms of action of GM-CSFR in sensory neurons are largely unknown but in hematopoietic cells, activation of GM-CSFR is known to stimulate cell signaling pathways regulating gene expression, including the JAK-STAT pathway (Janus kinase, JAK; signal transducer and activator of transcription protein, STAT) (Stosser, et al.,2011). Unfolding of JAK activity leads to dimerization and translocation of STAT family transcription factors to cell nucleus to regulate gene expression (Fortin, et al.,2007). The main aim of the present study was to identify molecules involved in GM-CSF-mediated signaling pathway in nociceptors and test their relevance to GM-CSF-induced pain.

Ion channels are the basis of sensory neuronal excitability and were previously suggested as molecular targets of GM-CSF signaling pathway (Bali, et al.,2013). Our preliminary screening (see below) revealed that GM-CSF selectively increased expression in DRG neurons of three voltage-gated sodium channels, Nav1.7, Nav1.8 and Nav1.9. Due to the primary role of these channels in the ability of DRG neurons to generate action potentials (APs), we hypothesized that GM-CSF might promote pain and hyperalgesia by acting on voltage-gated Na^+^ channels in nociceptors.

At least five different voltage-gated sodium channels are reportedly expressed in DRG, including the TTX-sensitive Nav1.1, Nav1.6 and Nav1.7 and the TTX-resistant Nav1.8 and Nav1.9 (Cummins, et al.,2000). Nav1.7, Nav1.8 and Nav1.9 channels are mainly distributed in small diameter DRG neurons which are related to pain initiation. NaV1.7 produces a rapidly-activating, rapidly-inactivating and slowly repriming current, contributing to the generation and propagation of action potentials and acting as a threshold channel regulating excitability (Francois-Moutal, et al.,2018)(Li, et al.,2018). Gain-of-function mutations within the Nav1.7 gene *SCN9A* lead to inherited pain disorders, such as erythromelalgia (IEM) and paroxysmal extreme pain disorder (PEPD). In contrast, loss-of-function *SCN9A* mutations result in congenital insensitivity to pain (CIP) (Cheng, et al.,2011; Dib-Hajj, et al.,2008; Jarecki, et al.,2010; Klein, et al.,2013; Sawal, et al.,2016). Nav1.8 mediates a slowly-inactivating sodium currents acting as a key component of the upstroke of the action potential and thus influences neuronal excitability and pain transmission. Mutation of Nav1.8 gene, *SCN10A*, is found in patients with peripheral neuropathy (Choi, et al.,2007; Lai, et al.,2002). The Nav1.9 channel has a slow kinetics and is responsible for persistent Na^+^ currents in nociceptors; together with the Nav1.7 it acts as a threshold channel for AP firing; it amplifies sub-threshold stimuli leading to AP bursts (REF). Gain-of-function mutations of Nav1.9 channel gene, *SCN11A* cause familial episodic pain syndrome (Huang, et al.,2014; Huang, et al.,2017).

In this study, we found GM-CSF significantly increase the excitability of DRG neurons in parallel with the increase the current density, mRNA and protein expression of Nav1.7, Nav1.8 and Nav1.9 channels. GM-CSF induces pain behaviors likely through up-regulation of Nav1.7, Nav1.8 and Nav1.9 channels by activating Jak2-Stat3 signal transduction pathway regulating Na^+^ channel expression.

## Results

### GM-CSF plays a crucial role in bone metastases cancer pain

Osteosarcoma is the most common form of primary bone cancer, in which pain is the most common symptom and is seen in 85% of patients (Yoneda, et al.,2015). Osteochondroma is the most common benign bone tumor and the majority of osteochondromas form an asymptomatic hard immobile painless palpable mass(de Mooij, et al.,2012). We first collected tumor tissue from osteosarcama and osteochondroma patients after surgery, and compared the expression levels of GM-CSF in these two tumor tissue types. The H&E staining show typical characteristics of a cancerous (Fig. 1A, low left) and a benign (Fig. 1A upper left) tissue of bones. Interestingly, osteosarcoma biopsy samples demonstrated a dramatically higher levels of GM-CSF than were seen in osteochondroma samples in immunohistochemistry study (Fig. 1A right panels; summarized in Fig. 1B, n = 8, *P < 0.05).

**Fig. 1:**
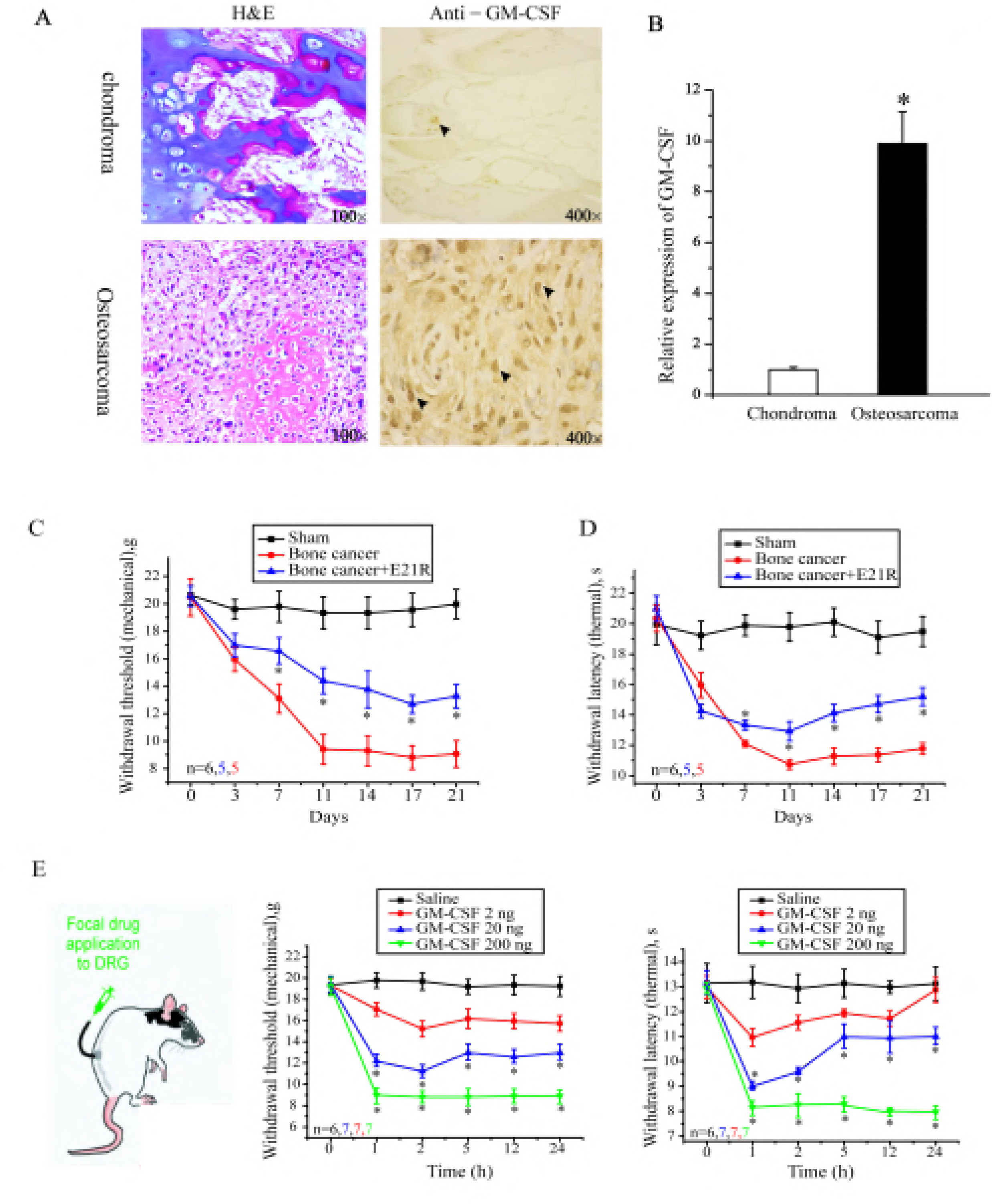
The role of GM-CSF in bone metastases cancer pain. A. High expression level of GM-CSF in osteosarcoma tissue sample. H&E staining of chondroma and osteosarcoma is shown on the left; immunohistochemical staining for GM-CSF indicated by arrows in chondroma and osteosarcoma is shown on the right. B. Summary results for immunohistochemical staining for GM-CSF (n = 8 per group, *P < 0.05). C and D. Effect of GM-CSF analogue GM-CSF (E21R, 25 μg/μl,3 μl), a competitive antagonist of GM-CSF on mechanical and thermal nociceptive responses in bone cancer pain model of rats. The mechanical paw withdrawal threshold and thermal paw withdrawal latency were measured at 3, 7, 11, 14, 17, and 21 days for the control group (black line + squares), the bone cancer group (red line + circles), and the bone cancer + E21R group (blue line + triangles). E. Dose-dependent effects of GM-CSF on paw withdrawal threshold to mechanical stimulus and on paw withdrawal latency to noxious heat at 1 h, 2 h, 5 h, 12 h and 24 h following focal DRG application via DRG cannula. Number of experiments is indicated as *n* in each panel. \**P* < 0.05 as compared to the vehicle saline.

To explore the role of GM-CSF in bone cancer pain, we first established the bone metastases cancer pain model induced by the implanted Walker 256 carcinoma cells. Consistent with previous reports(Wang, et al.,2011), mice with tumors in the tibia bone displayed a significantly lower withdrawal threshold for mechanical stimuli and a shortened withdrawal latency for thermal stimuli at 3-21 days after tumor cell injection (Fig. 1C, 1D). To assess the role of GM-CSF in these behavioral manifestations of pain hypersensitivity, we injected a mutant GM-CSF peptide with the glutamate-to-arginine substitution at the position 21 of a rat GM-CSF peptide sequence (E21R). E21R acts as a competitive antagonist of GM-CSF and can neutralize some of its biologic actions (Iversen, et al., 1996), however its effect on GM-CSF-mediated pain has not been tested before. E21R was injected in the vicinity of the tibia bone where the cancer cells had been implanted (see Methods). Treatment of the bone cancer rats with E21R significantly alleviated both mechanical and thermal hyperalgesia (Fig. 1C and 1D). The reduction of hyperalgesia was registered starting from day 7 after the establishment of the bone cancer model and lasted for the dyration of observation (21 days). These results indicate that GM-CSF is indeed involved in the bone cancer pain development, which is in agreement with a previous report that the bone cancer pain was attenuated following by a specific knockdown GM-CSF receptors in L4-L5 DRG of mice(Schweizerhof, et al.,2009).

To further attest that GM-CSF is pro-algesic and the primary afferent sensory nerve are the targeted sites of GM-CSF action, we evaluated the effect of direct focal GM-CSF infusion into the L5 DRG on pain-related behavior in naïve rats (Fig. 1E). Compared with the vehicle-treated rats, GM-CSF induced significant dose-dependent thermal and mechanical hyperalgesia which persisted for at least 24 hrs after injection. Indeed, focal injection of GM-CSF (20 - 200 ng) via the DRG cannula significantly increased sensitivity of rats to thermal and mechanical stimuli as measured with the Hargreaves and Von Frey tests (*P < 0.05). The nociceptive responses in rats received 2 ng of GM-CSF also showed a tendency towards sensitization but these effects not reach statistical significance.

### GM-CSF increase the excitability of small-sized DRG neuron

The fact that GM-CSF enhanced pain sensitivity when injected into DRG suggest that GM-CSF might directly sensitize nociceptors by increasing their excitability. To test this hypothesis, we performed current clamp recordings from the cultured DRG neurons in control conditions and after 24 hrs treatment with GM-CSF (200 ng/ml). For this study, small diameter (< 25 μM) DRG neurons were selected as these are predominantly nociceptors (Zheng, et al.,2013). AP firing was induced by trains of depolarizing current step from +50 to +500 pA injected with a 50 pA increment. GM-CSF significantly increased the number of APs induced by the depolarizing current pulses from 100 to 500 pA (Fig. 2A, B). GM-CSF also significantly lowered the rheobase currents (depolarization current threshold (CT) for eliciting the 1^st^ action potential) from 198.6 ± 9.8 pA (n = 63) to 83.1 ± 2.5 pA (n = 39, *P < 0.05); the action potential threshold voltage (TP) was also significantly decreased from −18.7 ± 0.5 mV (n = 23) to −20.4 ± 0.6 mV (n = 22, *P < 0.05). However, the resting membrane potential was not significantly changed (Fig. 2C, 1D). We also measured the effect of GM-CSF treatment on other properties of evoked APs such as AP amplitude (mV) and rate of depolarization (V/s), which are summarized in Table 1. Together, the above data indicate that GM-CSF increases intrinsic neuronal excitability of primary sensor neurons associated with pain.

**Table1.** Summarized effects of GM-CSF on parameters of action potential

| | Rhobase<br>currents(pA) | TP<br>(mV) | AP<br>amplitude<br>(mV) | Depolariz-<br>ation rate<br>(v/s) | AHP<br>amplitude<br>(mV) | AHP<br>duration<br>(ms) | RMP<br>(mV) | M-type K<sup>+</sup><br>current<br>(pA/pF) | Rin<br>(MΩ) |
| --- | --- | --- | --- | --- | --- | --- | --- | --- | --- |
| Control | 198.6±9.8<br>(63) | -18.6±0.5<br>(23) | 107.9±2.2<br>(19) | 16.2±0.6<br>(19) | 21.3±0.8<br>(23) | 19.4±0.8<br>(18) | 55.1±0.9<br>(n=43) | 3.93±1.0<br>(9) | 564.0±<br>51.9<br>(32) |
| GM-CSF | 83.1±2.5**<br>(39) | -20.4±0.6*<br>(22) | 113.5±1.8<br>(22) | 19.3±0.4*<br>(21) | 21.6±0.8<br>(26) | 21.6±1.3<br>(18) | 55.4±0.9<br>(58) | 3.54±0.9<br>(8) | 504.5±<br>24.1<br>(25) |

Rhobase currents: the depolarized current threshold for evoking the 1st action potential; TP: threshold potential; AP: action potential. RMP: resting membrane potential; Rin: input resistance.
\*p<0.05, \*\*p<0.01 compared with the control

**Fig. 2:**
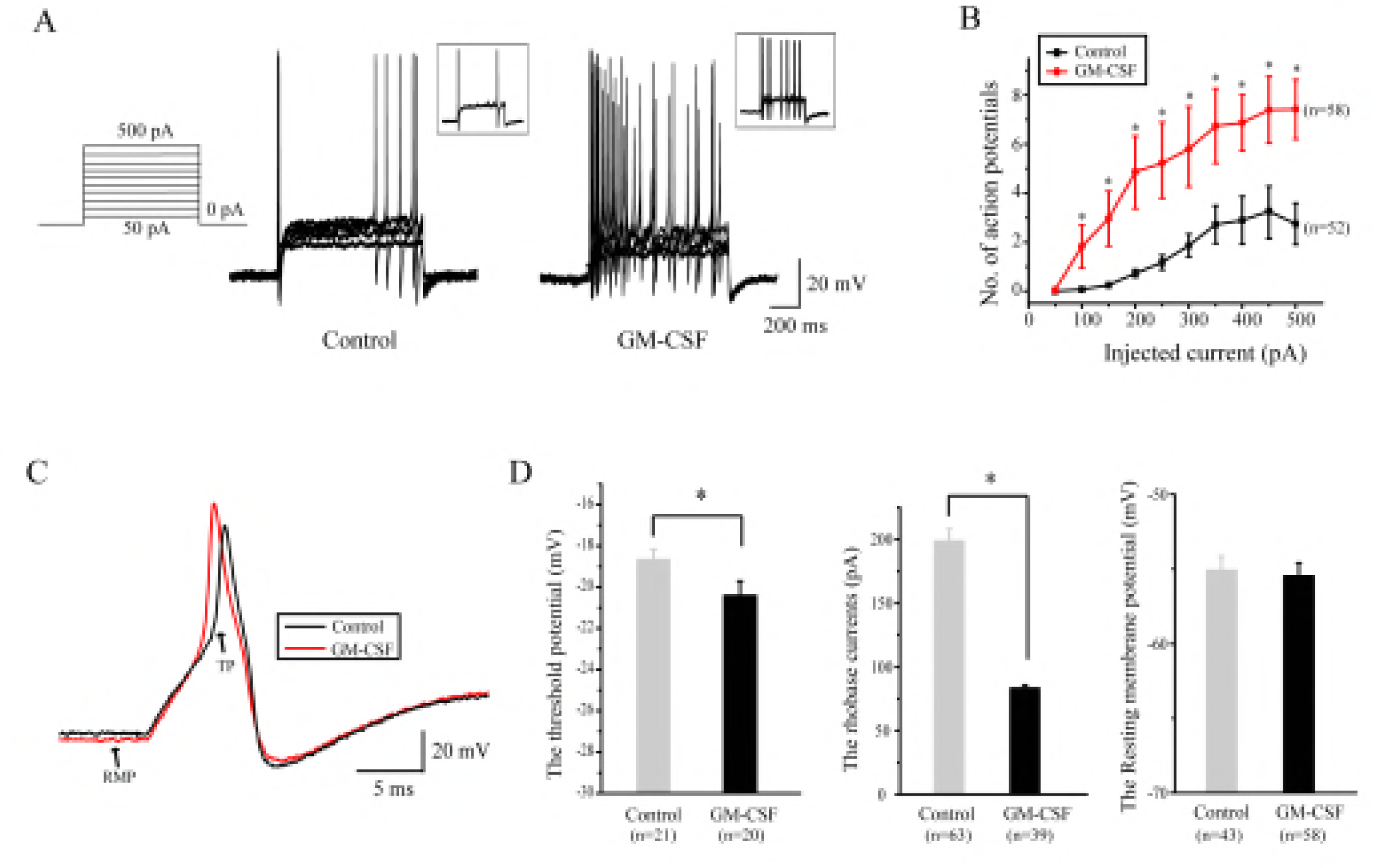
The effect of GM-CSF on the excitability of small-sized DRG neurons. A. Representatives of action potentials evoked by depolarizing current pulse (left), recorded from small-sized DRG neurons. B. Summary results for the effect of GM-CSF on numbers of action potential induced by increasing amplitudes of depolarizing currents. C. Single action potentials from A with expanded time scales. TP, threshold potential; RMP, rest membrane potential. D. Summary results for the effect of GM-CSF on the threshold potential, rhobase current and resting membrane potential (\**p*<0.05).

### GM-CSF increases activity and expression level of Nav1.7 Nav1.8 and Nav1.9 sodium channels

The sensitizing effect of a single focal in vivo injection of GM-CSF was long-lasting (Fig. 1E), in addition, intracellular action of GM-CSFRs has long been linked to transcriptional effects via the activation of JAK-STAT pathway. Thus, we hypothesized that the sensitizing effect of this growth factor might be mediated by changes in the expression of some intrinsic regulator(s) of excitability. Thus, to further explore the mechanisms for GM-CSF-induced hyperactivity of DRG nociceptors, we screened the effect of GM-CSF on ion channels which have been implicated in modulation of resting excitability of DRG neurons (Bernier, et al.,2018; Liu, et al.,2010; Zhang, et al.,2018; Zheng, et al.,2013). We tested the effect of treatment of cultured DRG neurons with 200 ng/μl GM-CSF (24 hrs) on the mRNA abundance of the following ion channel genes: *Scn9a* (Nav1.7), *Scn10a* (Nav1.8), *Scn9a* (Nav1.9), *Kcnd2* (Kv4.2), *Ano1* (TMEM16A), *P2rx3* (P2X3), *Kcng2* (Kv7.2), *Kcnq3* (Kv7.3). Among the transcripts tested, only the mRNAs of voltage-gated sodium channels Nav1.7, Nav1.8 and Nv1.9 were elevated (Supplementary Fig. 1). Next, we examined the effects of GM-CSF treatment on the current density of voltage-gated sodium currents in DRG neurons. Total, TTX-sensitive (TTX-S) and TTX-resistant Na^+^ currents were recorded (see Method) using whole-cell patch clamp. After pretreatment of DRG neurons with GM-CSF (200 ng/ml; 24hrs), the peak current density of total Na^+^ currents was increased from −96.0 ± 8.8 (pA/pF) (n = 41) to −154.2 ± 10.9 (pA/pF) (n = 26, \**P* < 0.05); the peak current density of TTX-S currents (mainly Nav1.7 currents) was increased from −70.2 ± 4.6 (pA/pF) (n = 66) to −98.6 ± 13.1 (pA/pF) (n = 41, \**P* < 0.05); the peak current density of TTX-R currents was increased from −82.2 ± 4.9 (pA/pF) (n = 66) to −111.6 ± 5.7 (pA/pF) (n = 41, \**P* < 0.05) (Fig 3A). We noted that the sum of the TTX-R and TTX-S current amplitudes produced larger value as compared to the total Na^+^ current measured, thus, the pharmacological separation of the macroscopic sodium currents on the basis of the TTX sensitivity is not complete. Yet, we believe that the TTX-R current fraction is predominantly Nav1.8/Nav1.9 while TTX-S fraction is predominantly Nav1.7, despite the cross-contamination(Cummins, et al.,2000).

**Fig. 3:**
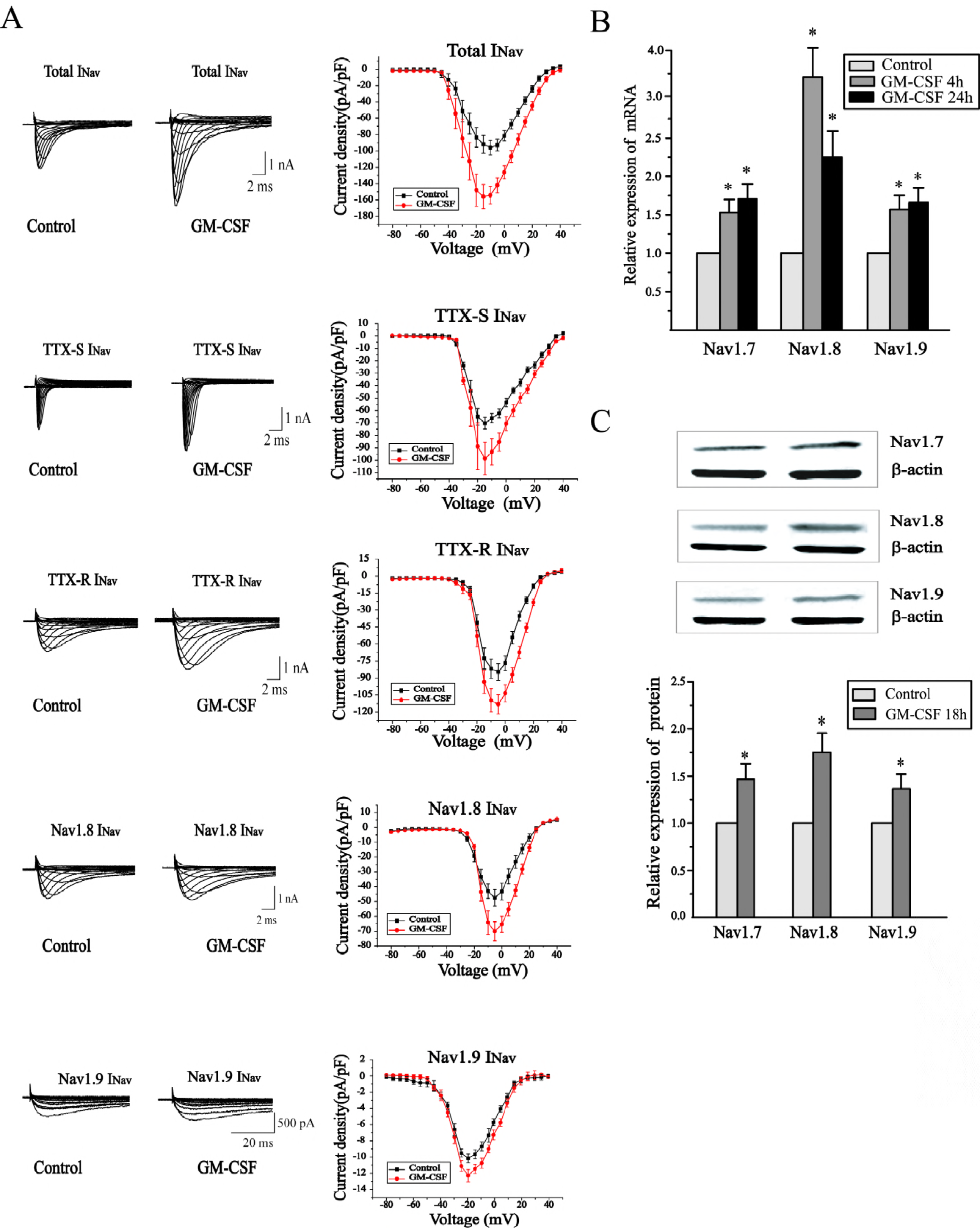
The effect of GM-CSF on the current amplitude and mRNA expression level of Nav1.7, Nav1.8 and Nav1.9 channels. A. Typical current traces and current desnity-voltage relationship of total TTX-S, TTX-R, Nav1.8 and Nav1.9 Na^+^ currents in cultured DRG cells after incubation with GM-CSF (200 ng /ml) for 24h. B. Relative mRNA expression of Nav1.7, Nav1.8 and Nav1.9 in cultured DRG cells after incubation of GM-CSF(200 ng /ml) 4h and 24h. C. Western blot analysis of expression levels of Nav1.7, Nav1.8 and Nav1.9 proteins in DRG neurons treated with GM-CSF (200 ng /ml) for 18h (n = 3, \**p*<0.05)

The TTX-R Na^+^ currents were further separated into Nav1.8-rich currents and Nav1.9-rich current fractions (see Method). After pretreatment of DRG neurons with GM-CSF, the peak current density of Nav1.8-rich current fraction was increased from −47.5 ± 5.5 (pA/pF) (n = 55) to −70.1 ± 6.4 (pA/pF) (n = 31, \**P* < 0.05); the Nav1.9-rich current fraction was increased from −9.3 (pA/pF) (n = 53) to −11.9 9 (pA/pF) (n = 30, \**P* < 0.05) (Fig. 3A). In sum, GM-CSF significantly increased the current amplitudes of nociceptor-specific Nav1.7 Nav1.8 and Nav1.9 currents.

We further measured mRNA levels of Nav1.7, Nav1.8 and Nav1.9 after the pre-treatment with GM-CSF at different time intervals. The mRNA levels of all three channels in DRG neurons were significantly increased after the GM-CSF pretreatment (200 ng/ml) for 4h and for 24h (Fig. 3B). Furthermore, the protein expression levels of Nav1.7, Nav1.8 and Nav1.9 channel in DRG neurons were also significantly increased after incubation with GM-CSF (200 ng/ml; 18 h; Fig. 3C).

### Down regulation of Nav1.7-Nav1.9 channels alleviate MG-CSF induced pain

To investigate whether the up-regulation of Nav1.7, Nav1.8 and Nav1.9 channels contributes to the GM-CSF-mediated pain we performed unilateral in vivo knockdown of either of the sodium channel subunit in rats using the anti-sense oligodeoxynucleotides (AS ODNs). AS ODNs against *Scn9a*, *Scn10a* and *Scn11a* (or a control mismatched ODN) were injected into the L5 DRG via the DRG cannula to offset the up-regulation of these Na^+^ channels, and then the effect of GM-CSF on pain behavior was tested. The efficiency knockdown was measured first; for this, the L5 DRGs was extracted following focal injection of saline, GM-CSF, ODN + GM-CSF, and then the mRNA expression levels of *Scn9a*, *Scn10a* and *Scn11a* were analized by quantitative PCR. In agreement with earlier data, the mRNA expression levels of *Scn9a*, *Scn10a* and *Scn11a* in DRG neurons were significantly increased after GM-CSF injection, and importantly, these increments were totally reversed by respective ODNs (Fig. 4A). Consistent with our earlier conclusion that up-regulation of Na1.7-Nav1.9 is a crucial factor in GM-CSF-induced hypersensitivity, ODNs against *Scn9a*, *Scn10a* and *Scn11a* significantly alleviated both mechanical and thermal hypersensitivity developed after the focal DRG application of GM-CSF(Fig. 4 B-D).

**Fig. 4:**
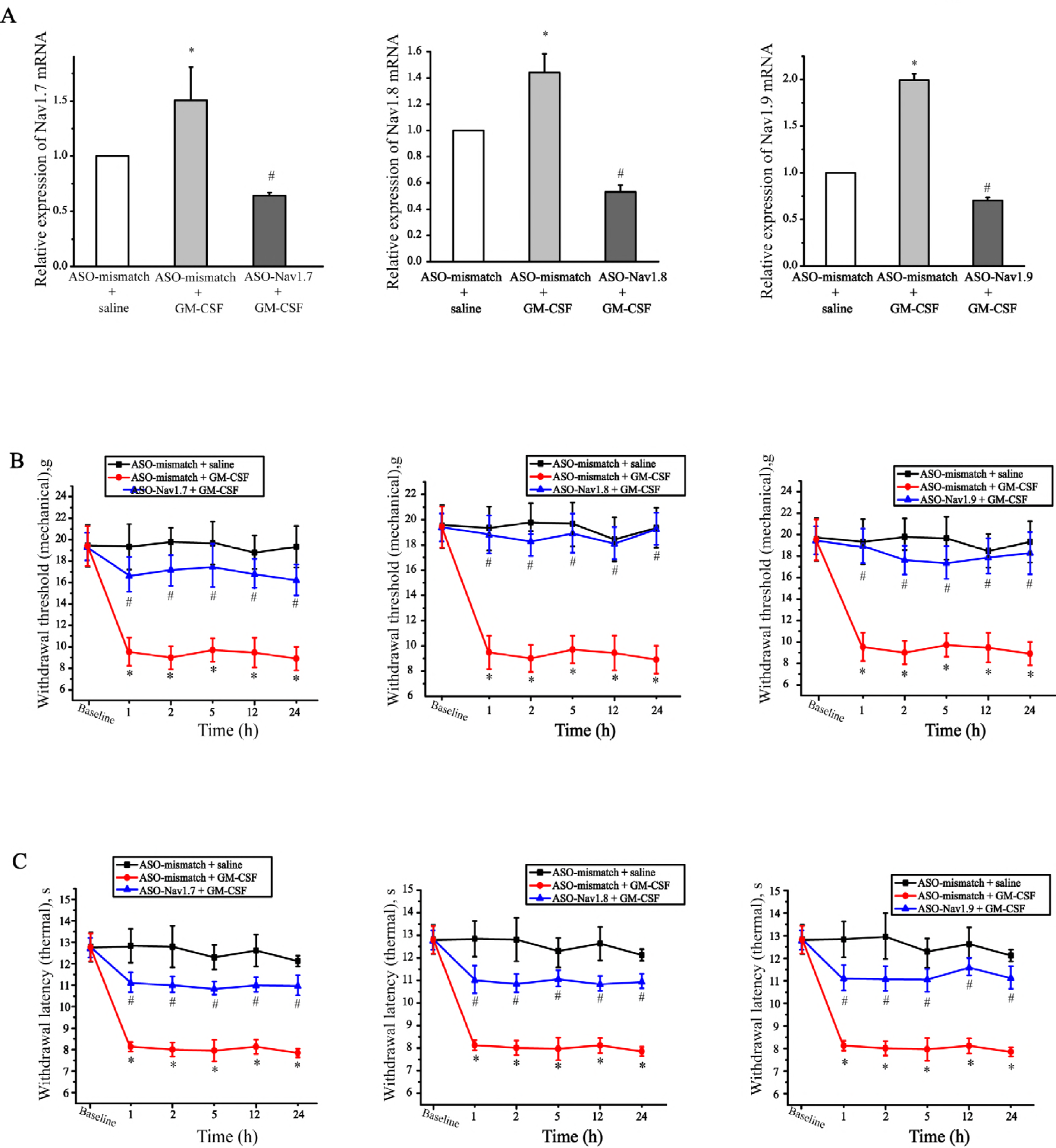
Down regulation of Nav1.7, Nav1.8 and Nav1.9 reverses nociceptive behavior evoked by GM-CSF. A. Application of antisense oligodeoxynucleotides (ASO) in DRG against Nav1.7, Nav1.8 and Nav1.9 (each ASO, 12.5 μg/rat, 5μl) significantly reduced the mRNA expression level of Nav1.7, Nav1.8 and Nav1.9 increased by GM-CSF treatment, and alleviated mechanical (B) and thermal hyperalgesia produced by the focal GM-CSF (200ng) application (C). (\**P* < 0.05 as compared to the vehicle saline; ^#^*P* < 0.05 with respect to the corresponding GM-CSF).

### Jak2-Stat3 signaling pathway is involved in GM-CSF induced up-regulation of Nav1.7-Nav1.9 channels

GM-CSF receptor is abundantly expressed in DRG (Schweizerhof, et al.,2009). Thus we hypothesized that the GM-CSF induced up-regulation of Nav1.7-Nav1.9 in DRG neurons could be mediated by the GM-CSF receptor and the related cellular signaling pathway. To test this, we focused on the Jak-Stat3/5 pathway since this is the key pathway for GM-CSF action in hematopoietic cells (Lilly, et al.,2001). Activated and phosphorylated states of Jak1, Jak2, Jak3, Stat3 and Stat5 were first measured in DRG neurons. As shown in Fig. 5A, after incubation of DRG cultures with GM-CSF for 25 min, phosphorylated Jak2 and Stat3 were significantly increased, but the level of phosphorylated Jak1 was not changed; phosphorylated Jak3 and Stat5 were not detected. These results indicate that GM-CSF is able to activate Jak2-Stat3 signaling pathway in DRG neurons. Is this activated Jak2-Stat3 signaling pathway responsible for GM-CSF–induced up-regulation of Nav1.7-Nav1.9 in DRG neurons? To test this, the acutely disassociated DRG neurons were incubated with GM-CSF with and without blockers of Jak2-Stat3 signaling pathway, AG490 (10 μM) and static (20 μM). Both compounds prevented up-regulation Nav1.7-Nav1.9 mRNA by GM-CSF (Fig 5B). In accord with these results, the GM-CSF-induced up-regulation of three Na^+^ channel proteins, measured using immunofluorescence method, were also prevented when the Jak2 inhibitor AG490 was present (Supplementary Fig. 2).

**Fig. 5:**
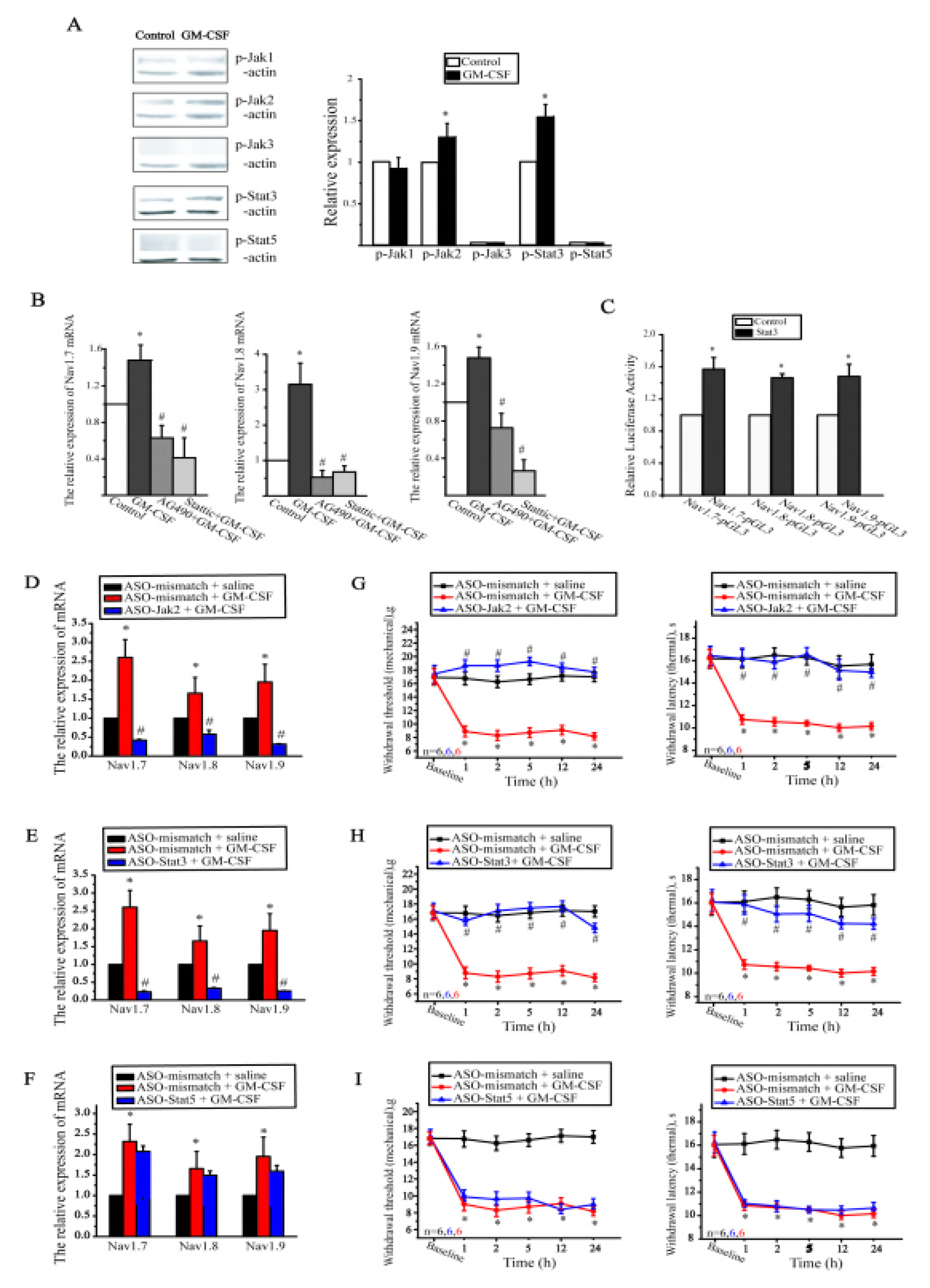
GM-CSF increase the mRNA expression level of Nav1.7, Nav1.8, Nav1.9 channel through Jak2-Stat3 signaling pathway. A. Relative expression of p-Jak1, p-Jak2, p-Jak3, p-stat3 and p-stat5 in DRG neurons after incubation with GM-CSF for 25 mins. B. Relative mRNA expression level of Nav1.7, Nav1.8, Nav1.9 in DRG neurons incubated with GM-CSF in the absence or presence of AG-490 (10 μM) and sttatic (20 μM) for 4h. C. Relative Luciferase activity in HEK 293 cells transfected with reporter vector containing Nav1.7, Nav1.8, Nav1.9 promoter regions (pGL3) co-expressed with either pcDNA3.1 (control) or pcDNA3.1-Stat3 cDNA. D-F. Relative mRNA level of Nav1.7, Nav1.8, Nav1.9 in ipsilateral DRGs (L5) of rats receiving anti-sense oligodeoxynucleotides (ASO) against different Jak and Stat signaling molecules (12.5 mg/rat, 5 μl) (n = 6 per group). D-I. Effect of ASOs against Jak and Stat signaling molecules (12.5 mg/rat, 5 μl) on hyperalgesia responses to mechanical and thermal stimuli induced by GM-CSF. ASOs were given through the DRG cannula for 4 days and then GM-CSF (200ng) was given. \**P* < 0.05 as compared to the vehicle saline; ^#^*P* < 0.05 with respect to the corresponding GM-CSF

As Stat3 was previously demonstrated to function as a transcriptional activator (Sharma, et al.,2018), we designed a luciferase reporter assay to determine if Stat3 is also act to regulate Nav1.7-Nav1.9 transcription. *Scn9a-Scn11a* promoter regions (relative to the transcription start site) were cloned into a luciferase reporter vector (pGL3 Basic plasmid) such that luciferase expression is driven by the *Scn9a-Scn11a* promoter sequences. We transfected these DNA constructs and either a control pcDNA3.1 plasmid or a pcDNA3.1-stat3 plasmid into HEK293 cells and measured the resulting luciferase activity. Luciferase activity in cells co-expressed with Scn9a, *Scn9a* or *Scn11a* promoter fragments and Stat3 was 1.57 ± 0.14 (n = 4, *P < 0.05), 1.5 ± 0.05 (n = 4, *P < 0.05), 1.5 ± 0.15 (n = 4, *P < 0.05) folds higher than that in cells co-expressed with Scn9a, *Scn9a* or *Scn11a* promoter fragments and the control pcDNA3.1 plasmid (Fig. 5C). These results implicate an important role for Stat3 in promoting Nav1.7, Nav1.8, and Nav1.9 gene expression.

Finally we assessed whether down regulation of Jak2-Stat3 signaling pathway would inhibit GM-CSF-induced up-regulation of Nav1.7-Nav1.9 and subsequently the pain behaviors. For this, AS ODNs against Jak2 and Stat3 were injected in DRG (L5) via DRG cannula as described above. AS ODNs against Jak 2 (Fig. 5D) and Stat3 (Fig. 5E) but not against stat5 (Fig. 5F) reversed the GM-CSF-induced up-regulation of Scn9a-*Scn11a* mRNA levels. Consistent with these results, AS ODNs against Jak 2 (Fig. 5G) and Stat3 (Fig. 5H) but not AS ODNs against stat 5 (Fig. 5I) prevented the development of the mechanical and thermal hyperalgesia produced by focal in vivo application of GM-CSF via the DRG cannula. Taken together, Jak2-Stat3 is a key target signaling pathway involved in GM-CSF induced pain behavior through up-regulation of Nav1.7-Nav1.9 channels.

## Discussion

In this study we demonstrate that GM-CSF promotes bone cancer-associated pain by enhancing excitability of bone afferents via the Jak2-Stat3-mediated upregulation of expression of nociceptor-specific voltage-gated sodium channels. First, we show that GM-CSF is highly expressed in osteosarcoma biopsy samples from human patients Second, we demonstrate that the competitive antagonist of GM-CSF, GM-CSF(E21R) is able to reduce both thermal and mechanical hyperalgesia in a rat model of bone cancer. Third, we show that GM-CSF increases excitability of peripheral nociceptors by upregulating functional expression of nociceptor-specific Na^+^ cannels, Nav1.7-Nav1.9. Using unilateral in vivo gene knockdown we further demonstrate that Na^+^ channel upregulation is indeed a necessary step in GM-CSF induced pain in vivo. Finally, using a set of genetic manipulations and assays, we delineated a molecular mechanism for GM-CSF induced initiation of pain in bone cancer: up-regulation of functional Nav1.7, Nav1.8 and Nav1.9 channel activity though the Jak2-Stat3 mediated activation of *Scn9a*, *Scn10a* and *Scn11a* gene transcription.

Several recent studies implicated contribution of GM-CSF to different types of pain, including cancer pain, neuropathic, inflammatory and osteoarthritic pain. (Cook, et al.,2013; Cook, et al.,2012; Nicol, et al.,2018). Yet, the exact mechanism and main molecular steps of the pro-algesic action of GM-CSF remained elusive. Our study fills this gap providing a full mechanistic concept for the effect.

Increased excitability of nociceptive neurons is a fundamental mechanism for pain. In turn, changes in excitability are ultimately linked to altered ion channel activity, thus, in this study we focused on ion channels in DRG neurons. All results pinpoint Nav.7-Nav1.9 channels as key determinants of the GM-CSF proalgesic action. 1) GM-CSF increased levels of *Scn9a*, *Scn10a* and *Scn11a* but not the other ion channels tested. 2) Consistent with above results, protein level of Nav1.7-Nav1.9 and the appropriate Na^+^ current fractions in nociceptive DRG neurons were also increased by GM-CSF. 3) Changes of DRG neuron excitability induced by GM-CSF were consistent with elevated Na^+^ channel activity: lowered rheobase, lowered threshold potential, but no significant change in resting membrane potential. Notably, GM-CSF did not change amplitude of M-type K^+^ current, which is another type of ion channels, important for setting resting excitability parameters of a neuron. 4) Down regulation of Nav1.7-Nav1.9 with anti-sense oligodeoxynucleotides alleviated GM-CSF-induced pain behavior. Although the last evidence does not directly prove involvement of these Nav channels since down regulation of them will probably inhibit any type of pain behavior anyway, the combined evidence implicate Nav channel mechanism as the most plausible and straightforward explanation for GM-CSF induced pain nonetheless.

We provide evidence that Jak2-Stat3 signaling pathway contributes to GM-CSF mediated up-regulation of Nav channels described above and, thus, to hyperalgesia associated with high GM-CSF levels, e.g. bone cancer. Activation of Jak by GM-CSF leads to the activation of the Stat family transcription factors, which dimerize and translocate to the nucleus upon activation and modulate gene expression (Choi, et al.,2011). In hematopoietic cells, GM-CSF exerts its biological functions mainly through activation of Jak2, which then activates Stat3 and Stat5 but not Stat2, Stat4 or Stat6 (Zgheib, et al.,2013). However, the signal transduction pathways mediated by GM-CSF and receptors are cell-type specific and and may differ significantly (Valdembri, et al.,2002). In the present study we found that in DRG neurons Jak2 and Stat3 are selectively phosphorylated following the GM-CSF treatment phosphorylated Jak1 was not affected and phosphorylated Jak3 and Stat5 were not fund at all. Consistent with these results, very low levels of Jak3 and Stat5 mRNA in DRG neurons were retrieved using the iBrain big data platform. Thus, Jak2-Stat3 is the dominant signaling pathway for GM-CSF to exert its function in DRG neurons.

Luciferase reporter assay provided strong evidence indicating that Stat3 is able to bind to the promoter regions of *Scn9a*, *Scn10a* and *Scn11a* genes to promote their transcription. In accordance with this observation, down regulation of Jak2 and Stat3 with anti-sense oligodeoxynucleotides reversed the GM-CSF induced elevation of mRNA expression level of these Nav channels. Importantly, these anti-sense oligodeoxynucleotides against *Scn9a-Scn11a* also alleviated the GM-CSF elicited pain behavior. These results not only describe a clear mechanismfor how Nav1.7-Nav1.9 channels are up-regulated by GM-CSF signaling pathway, but also indicate specific Jak-Stat pathway could be therapeutic target for pain treatment options..

GM-CSF is used clinically for treatment of myelodysplastic syndromes, aplastic anemia, tumor radiotherapy and chemotherapy-induced neutropenia(Garcia, et al.,2014). During these treatments, the most severe adverse reaction for GM-CSF is bone pain, with an incidence up to 90%(Stosser, et al.,2011). These clinical observations align very well with our results showing that GM-CSF induces pain behavior in rats when injected to DRG at a concentration of 20 ng/ml GM-CSF, which is lower than the blood concentration of ~600 ng/ml after a single-dose administration of GM-CAF in humans (Alexander,2016). Thus, clinically administered GM-CSF reaches sufficient blood concentrations to be able to sensitize bone periosteal nerves and nociceptive neurons through the mechanism described in this study.

In summary, in this study we provide mechanistic explanation for the role of GM-CSF in pain, specifically in pain associated with the bone cancer and with the therapeutic use of GM-CSF. This novel mechanism should be considered as a potential target for future pain treatments.

## Materials and methods

### Human subjects

The study was carried out in accordance with the ethical principles for medical research involving human subjects set out in the Helsinki Declaration, and was approved by the ethical committee at Hebei Medical University (Shijiazhuang, China). Osteosarcoma or chondroma tissues were obtained from 8 patients from the Fourth Hospital of Hebei Medical University. Each specimen was fixed with 4% paraformaldehyde for immunohistochemistry study. All patients or their relatives gave informed consent prior to their participation in the study.

### Animals

The animal protocols used in this study were approved by the Animal Care and Ethical Committee of Hebei Medical University under the International Association for the Study of Pain (IASP) guidelines for animal use. All surgeries were performed under sodium pentobarbital (Sigma) anesthesia, and all efforts were made to minimize animal suffering.

### Rat DRG neuron culture

Dorsal root ganglion (DRG) neurons were obtained from adult Sprague-Dawley rats (provided by Experimental Animal Center of Hebei Province) based on the protocol described previously (Du X, et al.,2014). Briefly, the ganglia were digested at 37°C with 1 mg/ml collagenase for 30 min. Ganglia were then mechanically triturated and washed twice with DMEM supplemented with 10% fetal calf serum. Thereafter, the DRG neurons were plated on poly-D-lysine-coated glass cover slips.

### Quantitative PCR

Total RNA was extracted using a commercial RNA isolation kit (RNAiso, Takara). Isolated RNA was dissolved in 20 μl DEPC-treated water and reverse-transcribed using an RT reagent kit (PrimeScript with gDNA Eraser, Takara) and a thermal cycler (Mastercycler, Eppendorf). Quantitative PCR reaction was performed using a kit (SYBR Premix Ex TaqII [Tli RNase H Plus], Takara), and the fluorescent DNA was detected and quantified with an FQD-48A(A4) system (BIOER). The PCR products were also run on a 2% agarose gel and were visualized using a gel imager (TFP-M/WL, Vilber Lourmat). For RT-PCR analysis, the following specific primers were used:

> Nav1.7-Forward: GCTCCAAGGACACAAAACGAAC, Nav1.7-Reverse: ATCAGACTCCCCAGGTGCAAT;
>
> Nav1.8-Forward: GACCCTTTCTACAGCACACACC, Nav1.8-Reverse: AAGTCCAGCCAGTTCCACG;
>
> Nav1.9-Forward: GCCCCTTCACTTCCGACT, Nav1.9-Reverse: GTCTTCCAGAGGCTTCGCTAC;
>
> GAPDH-Forward: CCAGCCTCGTCTCATAGACA, GAPDH-Reverse: CGCTCCTGGAAGATGGTGAT

### Luciferase reporter assay

The Stat3 luciferase reporter vector was designed to measure the binding of transcription factors to the enhancer, and was transfected into HEK293 cells with Lipofectamine2000 reagent (Invitrogen).

Fragments of rat *Scn9a*, *Scn10a* and *Scn11a* gene were amplified by PCR with following primers:

> Forward-GGCTCGAGAGCTTAAGGAAAGGAGGGTA, Reverse-GTAAGCTTTTTCCCCTTTGACTCCTTAC; corresponding to the promotor region of *Scn9a* (−286/+306).
>
> Forward-GGCTCGAGCCGTAGTAAGACCCTGCCTTG, Reverse-GTAAGCTTGAGACCCCAGCTCTGCAAAAC; corresponding to the promotor region of *Scn10a* (749/+124).
>
> Forward-GGCTCGAGCTTCACATGGTTGATCCATC Reverse-GTAAGCTTATTCTCGCTCTTGGCAGTA; corresponding to the promotor region of *Scn11a* (−51/+556 regions).

Amplified fragments were digested with appropriate restriction enzymes and cloned into pGL3 Basic (Promega). Luciferase activities were measured using a Dual Luciferase Assay Kit (Promega). Specific promoter activity was expressed as the relative activity ratio of firefly luciferase to Renilla luciferase.

### Western Blot

The DRG neuron lysates were prepared with RIPA lysis buffer. Equal amounts of protein were separated by sodium dodecyl sulfate polyacrylamide gel electrophoresis (SDS-PAGE) and electrotransferred to a polyvinylidene fluoride membrane. Membranes were blocked with 5% non-fat dairy milk and incubated with primary antibodies against Nav1.7 (1:500, Alomone), Nav1.8 (1:500, Abcam), Nav1.9 (1:200, Abcam), p-Jak1 (1:1000, Affinity), p-Jak2 (1:1000, Affinity), p-Jak3 (1:1000, Affinity), p-stat3 (1:1000, Epitomics), p-Stat5a (1:1000, Affinity) at 4°C overnight. This was followed by incubation with IRDye800-conjugated secondary antibody (1:20,000, Rockland) for 1 h at room temperature and subsequent scanning with the Odyssey Infrared Imaging System (LI-COR Biosciences). The integrated intensity for each detected band was determined using Odyssey Imager software (v3.0).

### Immunohistochemistry

Sections from human osteosarcoma or chondroma tissues were blocked with 0.3% hydrogen peroxide, followed by preincubation with 5% normal goat serum and then incubation with primary antibodies against GM-CSF (1:200, Pepro Tech) at 4 °C overnight. Next, the sections were incubated with the biotinylated secondary antibody, followed by streptavidin-horseradish peroxidase and diaminobenzidine, and then counterstained with hematoxylin. Staining intensities were determined by measurement of the integrated optical density (IOD) by light microscopy using a computer-based Image-Pro Morphometric System.

### Electrophysiology

Action potentials were recorded from dissociated rat small diameter DRG neurons (<25 μm) under current clamp using an Axopatch 200B amplifier and a Digidata 1322A converter (Axon Instruments). Pipettes (3–4 MΩ) were filled with solution containing (in mM) KCl 150, MgCl_2_ 5, HEPES 10, pH 7.4 adjusted with KOH. The bath solution contained (in mM): NaCl 160, KCl 2.5, MgCl_2_ 1, CaCl_2_ 2, glucose 10, HEPES 20 and pH 7.4 adjusted with NaOH. Small nociceptive DRG neurons were examined for evoked activity with a series of 1-s current injection from 0 pA to 500 pA in 50 pA increments. The rheobase currents was determined by the first action potential elicited by a series of depolarizing current injections that increased in 5 pA increments. The following values were measured in this study: resting membrane potential (RMP), threshold potential (TP), AP amplitude, depolarization rate (V/s).

Sodium currents were recorded from these small diameter DRG neurons under the voltage clamp mode in a whole-cell configuration. Pipettes (3–4 MΩ) were filled with solution containing (in mM): 70 CsCl, 30 NaCl, 30 TEA-Cl, 10 EGTA, 1 CaCl_2_, 2 MgCl_2_, 2 Na_2_ATP, 0.05 GTP, 10 HEPES, and 5 glucose, pH 7.3 with CsOH. The bath solution for DRG neurons was (in mM): 80 NaCl, 50 choline-Cl, 30 TEA-Cl, 2 CaCl_2_, 0.2 CdCl_2_, 10 HEPES, and 5 glucose, pH 7.3 with NaOH. The acquisition rate was 10 kHz and signals were filtered at 2.5 kHz. Series resistances were compensated by 60-80%. Currents were elicited by a 40 ms pulse from a holding potential of −120 mV to test potentials between −80 mV and +40 mV in 5 mV increments. The TTX-resistant (TTX-R) sodium currents (including both Nav1.9 and Nav1.8 currents) were recorded in the presence of 300 nm TTX in the external solution. The TTX-sensitive (TTX-S) sodium currents were obtained by digital subtraction of the TTX-R sodium currents from the total currents. Nav1.8 currents were then elicited by a prepulse of −70 mV for 500 ms before the test potentials from −80 mV to +40 mV in 5 mV increments in the same neuron. The Nav1.9 currents were obtained by digital subtraction of the Nav1.8 currents from the TTX-R sodium currents (based on protocols from (Qiu, et al.,2016)).

### Rat model of tumor-evoked pain

The Walker 256 carcinosarcoma breast cancer cells were provided by Shanghai Cell Bank of the Chinese Academy of Sciences. Wistar rats were injected intra-peritoneally with the Walker 256 cancer cells 0.5 ml (2 × 10^7^ cells/ml and 6–7 days later ascitic fluid was extracted. Sprague Dawley rats (180-200 g) were anesthetized by intraperitoneal injection of sodium pentobarbital (60-80 mg/kg). The right leg was shaved and the skin was disinfected with 70% (v/v) ethanol. A 1-cm-long rostro-caudal incision was made in the skin over the lower one-third of the tibia for easy exposure with minimal damage to muscles and nerves. The medullary canal was approached by inserting a 23-gauge needle proximally through a hole drilled in the tibia. The needle was then replaced with a 10-μl microinjection syringe containing the cells to be injected. A 5-μl volume of Walker 256 cells (4×10^6^/ml) or boiled cells (sham group) were injected into the bone cavity. After a 2-min delay to allow cells to fill the bone cavity, the syringe was removed and the hole was sealed using bone wax In some experiments GM-CSF antagonist, E21R (25 μg/μl, 3 μl, Life Tein LLC) or vehicle was injected in the vicinity of the tibia bone. The wound was closed using 1-0 silk threads and dusted with penicillin powder. The rats were allowed unrestricted movement in their cages after recovery and their general condition was monitored during the experiment.

### Focal application of drugs to DRG in vivo

All surgical procedures were performed under deep anesthesia with an i.p. injection of pentobarbital sodium (60-80 mg/kg). A DRG cannula for focal application of substances to DRG was implanted as previously described(Du X, et al.,2017). Briefly, a midline incision was made at the L4-L6 spinal level of adult male rats (Sprague Dawley; 180-200 g), and the L5 was identified at the midpoint of a link between both sides of the iliac crest. A 0.8-mm hole (~1 mm off the inferior edge of the transverse process) was drilled through the transverse process over the L5 DRG. Approaching of a ganglion was verified by the twitch of the paw. A hooked stainless steel blunt-tip cannula (inner diameter 0.64 mm, length 4 mm) was forced into the hole and connected to a polypropylene tube (inner diameter 0.41 mm, length 4.5 mm). The incision was closed with sutures, and the cannula was firmly fixed in place with dental cement. Intramuscular injection of benzylpenicillin (19 mg/0.1 ml) was given immediately after surgery. Postoperatively, rats were housed individually in plastic cages with sawdust flooring and supplied with water and food *ad libitum*. Animals were left to recover for at least 24 hours before the experiments were carried out. Animals developing signs of distress were humanely sacrificed.

### Antisense oligonucleotide knockdown

On the second day after DRG cannula implantation, rats were given through the cannula the antisense oligodeoxynucleotides (AS ODNs) against *Scn9a, Scn10a* or *Scn11a* (each at 12.5 μg in 5 μl) were given consecutively twice a day for 4 days. Mismatched ODNs were also given at matched time points. On the fifth day, mechanical and thermal sensitivity was assessed at 1 h, 2 h, 5 h, 12 h, and 24 h after the focal DRG application of GM-CSF (5 μl) or saline (5 μl), respectively. Five groups of animals were tested, injected as follows:

| Group | type of ODN pre-treatment | treatment type |
| --- | --- | --- |
| 1 | mismatched ODN | saline |
| 2 | mismatched ODN | GM-CSF |
| 3 | AS ODN *Scn9a* | GM-CSF |
| 4 | AS ODN *Scn10a* | GM-CSF |
| 5 | AS ODN *Scn11a* | GM-CSF |

> The following specific ASO were used:
>
> Mismatched ODN: TCACCCAGCACCCCCAACACATAGTT
>
> ASO- *Scn9a*: CTGGATCAACATGGTCTTCA
>
> ASO- *Scn10a*: CCAGAACCAAGCACAGAGGA
>
> ASO- *Scn11a*: CACCATCTGCATCATCATCA

### Mechanical hyperalgesia

Threshold sensitivity to mechanical stimuli was assessed using the von Frey method as described previously (Chaplan et al., 1994). Briefly, calibrated nylon filaments (Von Frey hair, Stoelting Co,) with different bending forces were applied to the midplantar surface of the right hind paw of the rats. The filaments were applied starting with the softest and continuing in ascending order of stiffness. A brisk withdrawal of the right hind limb was considered a positive response.

### Thermal hyperalgesia

The paw withdrawal latency in response to heat was tested using the Hargreaves method on the right hind paw of the rats using a radiant heat lamp source (Mengtai Technology Co., Ltd.). The intensity of the radiant heat stimulus was maintained at 20%. The time to withdrawal of the right hind paw (elapse time) was recorded.

### Statistics and analysis

All data are given as mean ± SEM. Differences between groups were assessed by paired or unpaired two-tailed Student’s *t* test. Comparisons of the behavioral data between groups at each individual time point were conducted using a two-way analysis of variance. Differences were considered significant at *P* ≤ 0.05. Statistical analyses were using OriginPro 9.1 (Originlab Corp.).

## Acknowledgments

This work is supported by the National Natural Science Foundation of China (91732108 to HZ), (31401199 to FZ) and (31571088 to XD); Key Basic Research Project of Applied Basic Research Program of Hebei Province (16967712D to XD); Biotechnology and Biological Sciences Research Council (Grants BB/R003068/1 and BB/R02104X/1 to NG). We greatly appreciate the collection of human osteosarcoma and chondroma tissues from Prof. Yueping Liu (The Fourth Hospital of Hebei Medical University, China).

## Author Contributions

YW, DZ, FX-Z and FZ performed the experiments and data analyses. HZ, FZ, XD and NG designed the experiments. HZ, FZ and NG wrote the paper. All authors have read, edited and approved the content of the manuscript.

